# mRNA-Based Designer APCs Elicit Robust CD8⁺ and CD4⁺ T Cell Responses

**DOI:** 10.1101/2025.09.16.676697

**Authors:** Toan Van Le, Shota Imai, Iriya Fujitsuka, Makie Ueda, Sayuri Nakamae, Shusaku Mizukami, Rikinari Hanayama, Tomoyoshi Yamano

**Affiliations:** Department of Immunology, Graduate School of Medical Sciences, Kanazawa University, Kanazawa, Japan; WPI Nano Life Science Institute (NanoLSI), Kanazawa University, Kanazawa, Japan; Department of Immune Regulation, SHIONOGI Global infectious Disease Division, Institute of Tropical Medicine, Nagasaki University, Nagasaki, Japan

**Author notes:** Correspondence to T.Y or R.H.

## Abstract

mRNA-based therapeutics have demonstrated notable success in SARS-CoV-2 vaccines and are emerging in cancer immunotherapy. However, conventional mRNA cancer vaccines are limited by the low immunogenicity of tumor-associated and neoantigens. We addressed this limitation by formulating a modular, liposome-based mRNA cocktail comprising three distinct mRNAs encoding tumor antigen, the co-stimulatory molecule CD80, and membrane-tethered IL-2. Administration of this mRNA mixture transforms somatic cells into ‘designer antigen-presenting cells (APCs)’ in vivo, which simultaneously express the antigen, a co-stimulatory molecule, and a cytokine. These designer APCs more effectively activated tumor antigen-specific CD8⁺ T cells than mRNA encoding the antigen alone and elicited robust anti-tumor immune responses. In addition, substituting IL-2 in the mRNA mixtures with membrane-tethered IL-12 led to the expansion and differentiation of endogenous antigen-specific Th1 helper T cells in vivo. Importantly, this platform activated NY-ESO-1-specific CD8⁺ T cells both in human PBMCs in vitro and in HLA-A*02:01-transgenic mice, highlighting its translational potential. This modular mRNA strategy reprograms somatic cells in situ into designer APCs, providing a flexible and translatable platform for precision immunotherapy.

## Introduction

mRNA-based therapeutics have dramatically accelerated the development of infectious disease vaccines. This accelerated timeline is evident in the significant progress achieved in less than a year for SARS-CoV-2, a process that typically takes several years^1, 2^. This advancement, highlighted by the practicality and safety of mRNA vaccines, has increased interest in mRNA therapeutics globally. As a result, the development of vaccines for various infectious diseases and cancer is being pursued^3, 4^.

In cancer immunotherapy, mRNA encoding cancer neoantigens is currently being advanced in clinical trials^4, 5^. Recent efforts have focused on vaccines that encode multiple sequences of neoantigens found in cancer cells^6, 7^. However, in the context of cancer vaccines, the effectiveness of mRNA on its own remains limited, often necessitating the concurrent use of immune checkpoint inhibitors and other anti-cancer treatments^8^. This highlights the need for more efficient methods to activate CD8^+^ T cells for enhanced anti-tumor responses.

The delivery of mRNA into cells triggers a cascade of events, starting with its translation into proteins, which are subsequently degraded by the proteasome. The resulting peptides are then complexed with major histocompatibility complex class I (MHC I) molecules and displayed on the cell surface for recognition by CD8^+^ T cells^9^. However, T cell activation is not solely dependent on T cell receptor (TCR) engagement; it also requires co-stimulatory signals^10^. While professional antigen-presenting cells (APCs) express these co-stimulatory molecules, non-APCs lack them and are unable to directly activate naïve CD8^+^ T cells upon mRNA uptake. In such cases, cross-presentation by professional APCs becomes essential^11^. Non-APCs that have taken up mRNA-encoded antigens rely on antigen transfer to dendritic cells, particularly XCR1⁺ conventional type 1 DCs (cDC1s), which are specialized for cross-presentation and subsequent activation of CD8⁺ T cells^12^. This dependency implies that the therapeutic potential of mRNA vaccines may be constrained by the efficiency of antigen handover and the functionality of cross-presenting DC subsets. To address this limitation, recent efforts have focused on the development of lipid nanoparticles (LNPs) that preferentially target dendritic cells, including both cDC1s and migratory cDC2s. By engineering ionizable lipids or formulating mRNA with DC-targeting delivery platforms, such as lipid nanoparticles (LNPs) or lipoplexes (LPX), it is now possible to selectively deliver mRNA into DCs. This targeted approach bypasses the need for antigen transfer from non-APCs and enables direct expression of antigen and co-stimulatory molecules within dendritic cells^13, 14^. These delivery systems have demonstrated enhanced CD8⁺ T cell priming and improved therapeutic efficacy in preclinical cancer models, reinforcing the rationale for APC-directed mRNA delivery as a key component of next-generation cancer vaccines.

In this study, we propose a novel approach to further enhance mRNA vaccine efficacy by introducing mRNA constructs that encode not only tumor antigens, but also co-stimulatory molecules and cytokines, thereby transiently reprogramming recipient cells into ‘designer APCs’. This strategy complements the conventional reliance on natural APC subsets by functionally reprogramming transfected cells to acquire key immunostimulatory features, thereby providing an alternative route for antigen presentation. These designer APCs are capable of directly activating CD8⁺ T cells and orchestrating a strong cytotoxic immune response. Moreover, substituting IL-2 with cytokines such as IL-12 enables the in vivo differentiation of antigen-specific Th1 helper T cells, highlighting the potential of this strategy to precisely modulate T cell differentiation in a context-dependent manner.

## Result

### Generation of designer APCs for activation of antigen-specific CD8⁺ T cells

To engineer designer APCs capable of activating antigen-specific CD8⁺ T cells, we synthesized a set of mRNAs encoding OVA-MITD (a model antigen containing OT-I and OT-II epitopes, fused to a minimal MHC class I trafficking domain (MITD to enhance antigen presentation)^15^, the costimulatory molecule CD80, and membrane-anchored IL-2 (achieved by fusing IL-2 to the extracellular domain of CD8α). Plasmids used for in vitro transcription were engineered to include a 128-nucleotide poly(A) tail for enhanced stability and incorporated N1-methylpseudouridine triphosphate (m¹ΨTP) in place of UTP (Fig.1a, Supplementary Fig.1a-c)^15, 16^. These mRNAs were transfected into MC38 cells, resulting in robust cell surface expression of MHC I, CD80, and IL-2, as confirmed by flow cytometry (Fig.1b, c). This indicated successful reprogramming of MC38 cells into ‘designer APCs’ simultaneously presenting antigen, co-stimulatory molecule, and cytokine. When co-cultured with OVA-specific CD8⁺ T cells (OT-I), MC38 cells expressing all three components, OVA-MITD, CD80, and IL-2, induced the most robust OT-I proliferation, compared to cells expressing OVA-MITD alone or in combination with either CD80 or IL-2 (Fig.1d, e, Supplementary Fig.1d, e).

**Fig 1.**
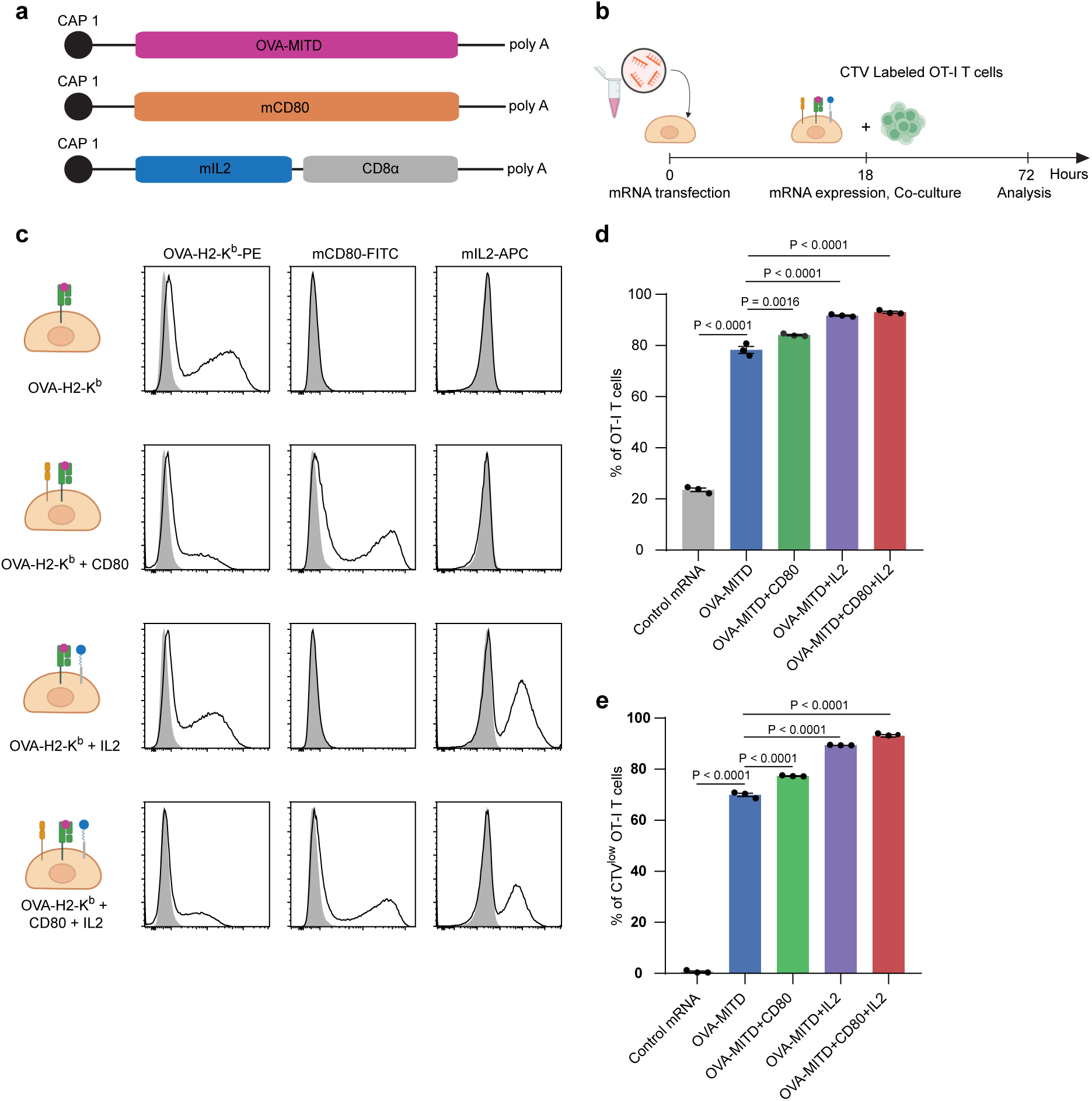
Generation of designer APCs for activation of antigen-specific CD8⁺ T cells. (a) Schematic of each mRNA construct encoding OVA-MITD, mouse CD80 or membrane-anchored mouse IL-2. 5′ cap, 5′ UTR, open reading frame, 3′ UTR and 128-nt poly(A) tail are indicated. (b) Co-culture scheme. MC38 cells were harvested 18 h after mRNA transfection, mitomycin C (MMC)-treated, and co-cultured with CellTrace Violet-labelled OT-I CD8⁺ T cells. (c) Flow-cytometric analysis of surface OVA-K^b^ complexes, CD80, and membrane-anchored IL-2 on MC38 cells 18 h after transfection. Filled gray histograms: mock-transfected MC38 cells; open black histograms: mRNA-transfected cells. Proliferation was analyzed 72 h later. (d) Frequencies of OT-I cells. (e) Frequencies of CTV^low^ OT-I cells. Data are mean ± SEM of technical triplicates. Statistical significance was determined by one-way ANOVA followed by Dunnett’s multiple-comparison test.

### In vivo generation of designer APCs stimulates antigen-specific CD8⁺ T cells

We examined whether in vivo generation of designer APCs could promote clonal expansion of endogenous antigen-specific CD8⁺ T cells. Mice were administered control mRNA, OVA-MITD mRNA alone, or a mixture of OVA-MITD, CD80, and IL-2 mRNAs once weekly for three consecutive weeks. All mRNAs were formulated using the in vivo-jetRNA⁺ liposomal delivery reagent for efficient systemic delivery. One week after the final injection, spleens were harvested and analyzed for OVA-specific CD8⁺ T cells using MHC class I tetramer (Fig.2a). Consistent with the in vitro results, OVA-tetramer⁺ CD8⁺ T cells expanded more robustly in the combination group (56%) than in the OVA-MITD group (24.3%) (Fig. 2b, c, Supplementary Fig. 2a). The expanded CD8⁺ T cells exhibited a CD44^hi^CD62L^low^ phenotype, characteristic of effector or effector memory T cells, and expressed IFN-γ (Fig.2d, Supplementary Fig.2b, c). Total splenocyte numbers were comparable across all groups (Supplementary Fig.2d), suggesting that the observed expansion of OVA-specific CD8⁺ T cells was not due to non-specific activation, but rather reflects a targeted immune response.

**Fig 2.**
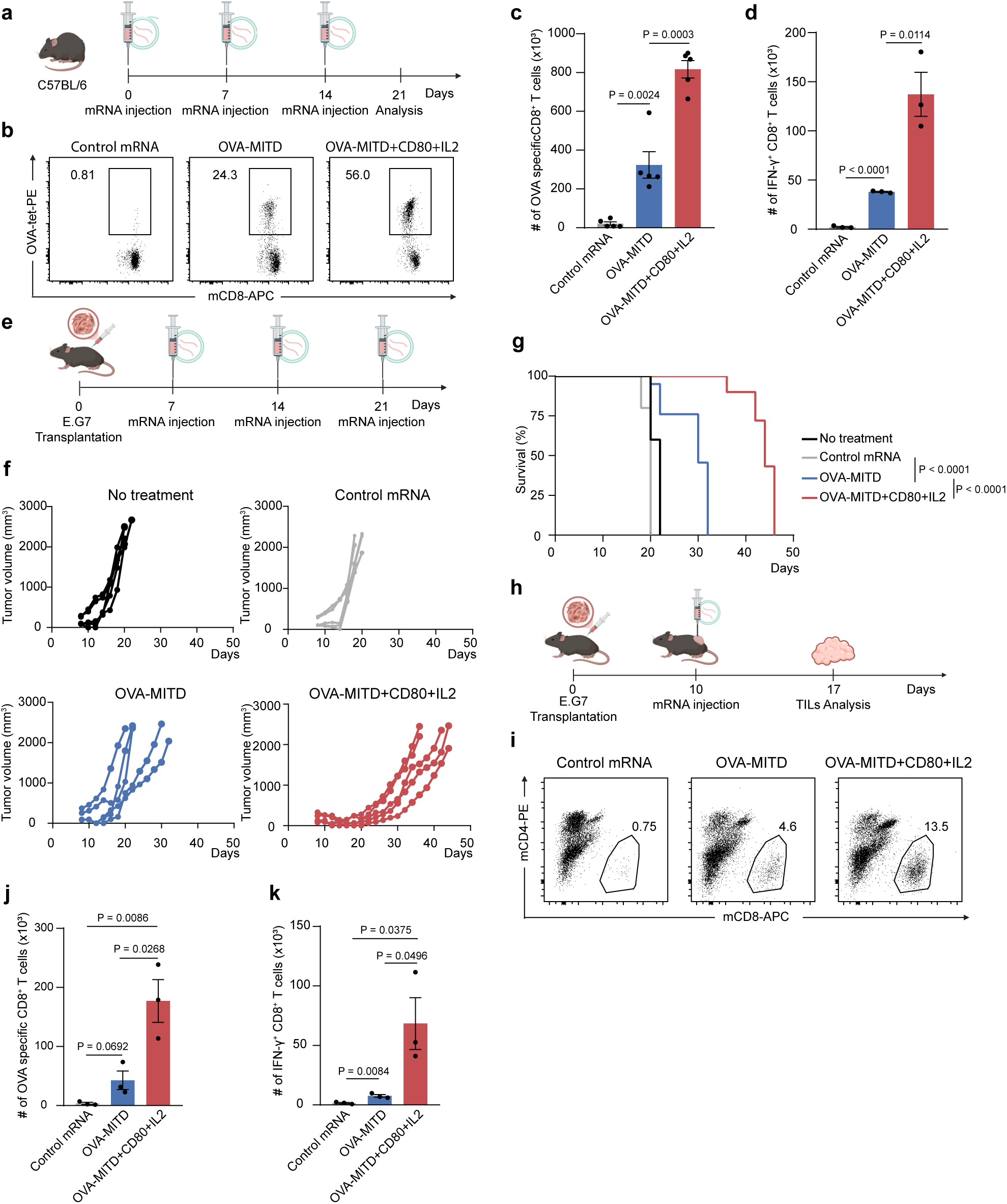
In vivo generation of designer APCs stimulates antigen-specific CD8⁺ T cells. (a) C57BL/6 mice received three weekly i.v. injections (days 0, 7, 14) of control mRNA, OVA-MITD mRNA (5 µg + 10 µg control), or OVA-MITD + CD80 + membrane-anchored IL-2 mRNAs (5 µg each; total 15 µg). Spleens were collected 7 d after the final dose. (b) Representative dot plots of OVA-specific CD8⁺ T cells. (c) Number of tetramer⁺ CD8⁺ T cells in the spleens. (d) Number of IFN-γ⁺ CD8⁺ T cells. (e) Mice were subcutaneously inoculated with OVA-expressing EL-4 cells (E.G7), followed by intravenous administration of mRNAs on days 7, 14, and 21. (f) Tumor growth curves are presented. (g) Kaplan-Meier survival curve. (h) E.G7 cells were subcutaneously implanted into C57BL/6 mice. When tumors reached ∼100 mm³, mice received intravenous injections of control mRNA, OVA-MITD mRNA alone, or a combination of mRNAs. Tumor-infiltrating lymphocytes were analyzed one week later. (i) Representative dot plots of TILs. (j) Number of tetramer⁺ CD8⁺ T cells. (k) Number of IFN-γ⁺ CD8⁺ T cells. Data are mean ± SEM. Statistical significance was assessed by using an unpaired, two-tailed Student’s t-test (c, d, j, k) and by the log-rank (Mantel-Cox) test for survival analysis (g).

To assess therapeutic efficacy, we employed a tumor model in which E.G7 cells (OVA-expressing EL-4 derivatives) were subcutaneously implanted into C57BL/6 mice. When tumors reached 100 mm³, mice received weekly intravenous injections of control mRNA, OVA-MITD mRNA alone or the combination mixture for three weeks (Fig.2e). While OVA-MITD mRNA monotherapy delayed tumor growth and extended survival relative to control, the combination treatment led to even greater tumor suppression and prolonged survival (Fig. 2f, g).

To elucidate the underlying mechanisms, we analyzed tumor-infiltrating lymphocytes one week after mRNA administration (Fig. 2h). CD8⁺ T cells constituted 4.6% of TILs in the OVA-MITD group, but increased to 13.5% in the combination group (Fig. 2i, j, Supplementary Fig.2e-g). Consistently, tumor weights were significantly lower in the combination group (Supplementary Fig.2h). Furthermore, a higher proportion of CD8⁺ T cells expressed IFN-γ in the combination group (Fig. 2k), indicating enhanced infiltration and activation of CD8⁺ T cells in the tumor microenvironment. These results demonstrate that in vivo-generated designer APCs expand antigen-specific CD8⁺ T cells and elicit potent anti-tumor immunity in the OVA antigen model.

### In vivo generation of designer APCs expands neoantigen-specific CD8⁺ T cells

Using Rpl18 as a model neoantigen in the MC38 tumor system, we investigated whether in vivo generation of designer APCs could expand neoantigen-specific CD8⁺ T cells. Mice were administered control mRNA, Rpl18-MITD mRNA alone, or a mixture of Rpl18-MITD, CD80, and IL-2 mRNAs once weekly for three consecutive weeks, in accordance with the administration protocol established in the OVA model (Fig.3a, Supplementary Fig.3a). One week after the final injection, spleens were harvested and analyzed for Rpl18-specific CD8⁺ T cells using MHC class I tetramers. Mice receiving Rpl18-MITD alone exhibited modest expansion of tetramer-positive CD8⁺ T cells, whereas the combination treatment induced robust expansion and differentiation into CD44^hi^CD62L^low^ effector and effector memory phenotypes (Fig.3b-d, Supplementary Fig.3b-d). As observed in the OVA model, total splenocyte numbers remained unchanged across groups (Supplementary Fig.3e), suggesting that the response was antigen-specific rather than due to non-specific activation.

**Fig 3.**
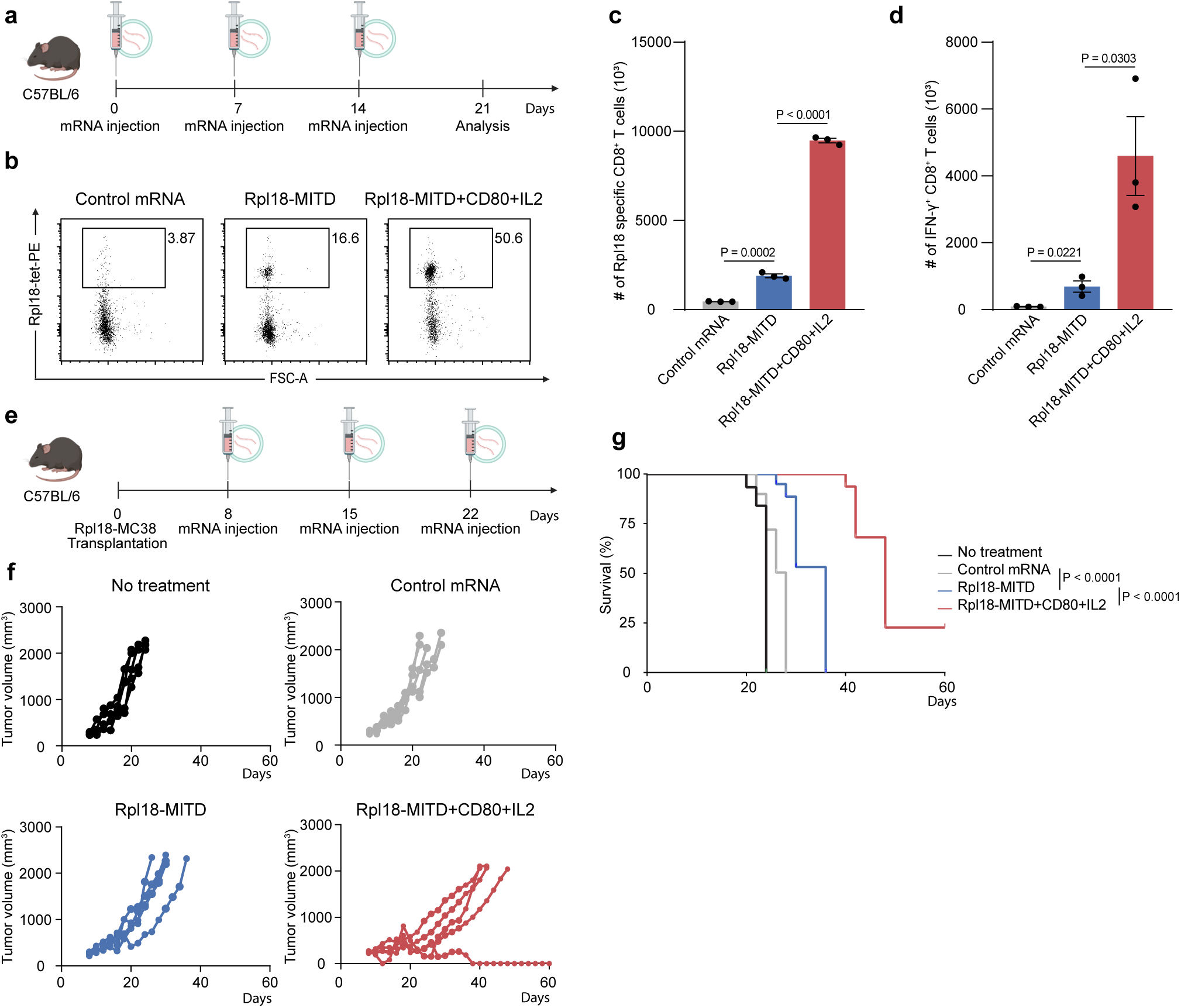
In vivo generation of designer APCs expands neoantigen-specific CD8⁺ T cells. (a) C57BL/6 mice received three weekly i.v. injections (days 0, 7, 14) of control mRNA (15 µg), Rpl18-MITD mRNA (5 µg) + control mRNA (10 µg), or Rpl18-MITD (5 µg)+ CD80(5 µg) + membrane-anchored IL-2(5 µg) mRNAs. Spleens were collected 7 d after the final dose. (b) Representative dot plots of Rpl18-specific CD8⁺ T cells in the spleens. (c) Number of tetramer⁺ CD8⁺ T cells in the spleens. (d) Number of IFN-γ⁺ CD8⁺ T cells in the spleens. (e) Mice were subcutaneously inoculated Rpl18-MC38 cells, followed by intravenous administration of mRNAs on days 7, 14, and 21. (f) Tumor growth curves are presented. (g) Kaplan-Meier survival curves of tumor-bearing mice. Data are mean ± SEM. Statistical significance for (c,d) was determined by using an unpaired, two-tailed Student’s t-test, and survival differences in (g) by the log-rank (Mantel-Cox) test.

To assess therapeutic efficacy, Rpl18-MC38 tumors were subcutaneously implanted and allowed to grow to approximately 100 mm³. Following the same weekly schedule as the OVA model, mice received either control mRNA, Rpl18-MITD alone or the combination mRNA mixture once per week for three weeks (Fig.3e). While Rpl18-MITD alone modestly delayed tumor growth, the combination treatment resulted in significantly greater tumor suppression and extended survival (Fig.3f, g). These results demonstrate that in vivo generation of designer APCs not only promotes the expansion and functional differentiation of neoantigen-specific CD8⁺ T cells, but also enhances anti-tumor immunity and improves survival compared to antigen-only mRNA vaccination.

### In vivo and in vitro evaluation of HLA-A*02:01-restricted responses induced by humanized designer APCs

To assess the function of designer APCs in an HLA-A*02:01-restricted humanized mouse model, we used HHD mice, which express a chimeric MHC class I molecule composed of the α1 and α2 domains of human HLA-A*02:01, the α3 domain of murine H-2Db, and human β2-microglobulin^18^. This model enables the evaluation of HLA-A*02:01-restricted CD8⁺ T cell responses in vivo. As a representative tumor antigen, we selected NY-ESO-1, a cancer/testis antigen widely expressed in various malignancies, and used its well-characterized HLA-A*02:01-restricted epitope (amino acids 157-165)^19^. Mice were administered either mRNA encoding NY-ESO-1-MITD alone or a combination of mRNAs encoding NY-ESO-1-MITD, CD80, and IL-2 (Fig.4a, Supplementary Fig.4a). NY-ESO-1-MITD alone failed to induce significant expansion of antigen-specific CD8⁺ T cells, whereas the combination treatment resulted in robust expansion and differentiation of CD44^hi^CD62L^low^ effector/memory CD8⁺ T cells (Fig.4b-d, Supplementary.4b-d). Consistent with earlier results, total splenocyte numbers remained unchanged across groups, indicating antigen-specific activation (Supplementary Fig.4e). To assess cytotoxic activity, we performed an in vivo killing assay using a 1:1 mixture of splenocytes pulsed with NY-ESO-1 peptide (CellTrace Violet-labeled) and control peptide (CFSE-labeled), injected into mRNA-treated mice seven days after the final injection (Fig.4e). NY-ESO-1 peptide-pulsed target cells were selectively eliminated in mice treated with the mRNA mixture, but not in those receiving control mRNA or NY-ESO-1-MITD mRNA alone (Fig.4f, g). These findings demonstrate that in vivo-generated designer APCs can effectively induce functional antigen-specific CTLs within an HLA-A*02:01-restricted setting.

**Fig 4.**
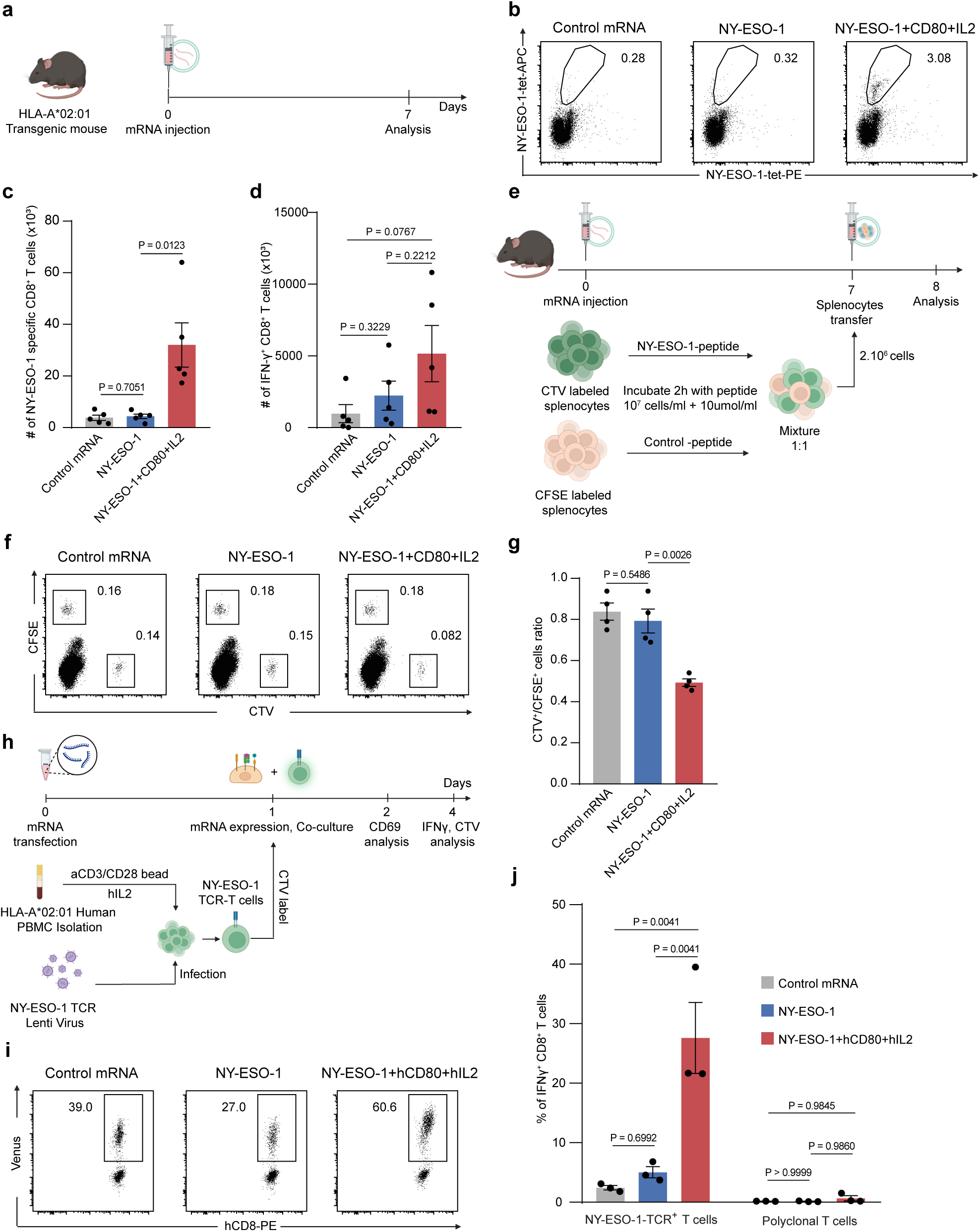
Generation of humanized designer APCs activates NY-ESO-1-specific CD8⁺ T cells. (a) HLA-A*02:01 transgenic mice received a single i.v. dose of control mRNA (15 µg), NY-ESO-1-MITD mRNA (5 µg) plus control mRNA (10 µg), or NY-ESO-1-MITD (5 µg) + mouse CD80 (5 µg) + membrane-anchored mouse IL-2 (5 µg) mRNAs. The spleens were collected 7 d later. (b) Representative dot plots of NY-ESO-1-specific CD8⁺ T cells in the spleens. (c) Number of tetramer⁺ CD8⁺ T cells in the spleens. (d) Number of IFN-γ⁺ CD8⁺ T cells in the spleen. (e) In vivo killing assay: on day 0 mice were treated as in (a). On day 7, 1 × 10⁶ NY-ESO-1-pulsed splenocytes (CTV-labelled) and 1 × 10⁶ control-peptide-pulsed splenocytes (CFSE-labelled) were transferred i.v.; 24 h later, spleens were harvested and target-cell killing was analysed by flow cytometry. (f) Representative dot plots of CTV⁺ and CFSE⁺ target cells. (g) Ratio of CTV⁺/CFSE⁺ cells in spleen. (h) Schematic of the in vitro co-culture assay. (i) Representative dot plots of NY-ESO-1-TCR (Venus) positive CD8 T cells. (j) IFN-γ expression in NYESO1-TCR⁺ and polyclonal T cells after 3 days of stimulation with HEK293 cells transfected with control mRNA, NYESO1-MITD mRNA, or a mixture of NYESO1-MITD, hCD80, and hIL-2 mRNAs. Statistical significance was determined using an unpaired, two-tailed Student’s t-test for panels (c), (d), and (g), and using two-way ANOVA followed by Tukey’s multiple comparisons test for panel (j).

Following in vivo evaluation in the HHD mouse model, we next examined the function of designer APCs in a fully human context by performing in vitro co-culture assays using TCR-transduced primary human PBMCs obtained from an HLA-A*02:01-positive donor. A NY-ESO-1-specific TCR was introduced into PBMCs derived from an HLA-A*02:01-positive healthy donor via lentiviral transduction^17^. HEK293T cells were then transfected with either control mRNA, NY-ESO-1-MITD mRNA alone, or a combination of NY-ESO-1-MITD, human CD80, and membrane-anchored human IL-2 mRNAs to generate designer APCs (Fig.4h, Supplementary Fig.4f, g). When co-cultured with the TCR-engineered T cells, only the designer APCs expressing all three components induced robust T cell activation, as demonstrated by the expansion of antigen-specific T cells and the production of IFN-γ (Fig.4i, j, Supplementary Fig.4h, i).

### In vivo generation of designer APCs induces Th1 differentiation of antigen-specific CD4⁺ T cells

To extend the functionality of designer APCs beyond CD8⁺ T cell activation, we engineered constructs to promote the differentiation of antigen-specific CD4⁺ helper T cells. We designed mRNAs encoding membrane-anchored IL-12 and a fusion protein comprising the OVA peptide fused to the MHC class II molecule (MHC-II) I-A^b^ via a flexible linker, enabling direct epitope presentation without intracellular processing (Fig.5a, Supplementary Fig.5a, b). Mice were intravenously injected with either OVA-MHCII mRNA alone or a combination of OVA-MHCII, CD80, and IL-12 mRNAs once weekly for two consecutive weeks. One week after the final injection, spleens were harvested and analyzed by flow cytometry using MHC-II tetramers to identify OVA-specific CD4⁺ T cells (Fig.5b). In mice treated with OVA-MHCII mRNA alone, only minimal expansion of antigen-specific CD4⁺ T cells was observed. In contrast, mice receiving the combination mRNA mixture exhibited a marked increase in the frequency of MHC-II tetramer⁺ CD4⁺ T cells (Fig. 5c, d, Supplementary Fig.5c, d). Among these cells, 27% expressed T-bet, a master regulator of Th1 differentiation^20^ (Fig.5e, f, Supplementary Fig.5e). Furthermore, upon ex vivo stimulation with OVA peptide, a substantial proportion of CD4⁺ T cells from the combination group produced IFN-γ, confirming functional Th1 polarization (Fig.5g, Supplementary Fig.5f). These results demonstrate that in vivo generation of designer APCs encoding an MHC-II-linked antigen and IL-12 effectively promotes the expansion and Th1 differentiation of antigen-specific CD4⁺ T cells.

**Fig 5.**
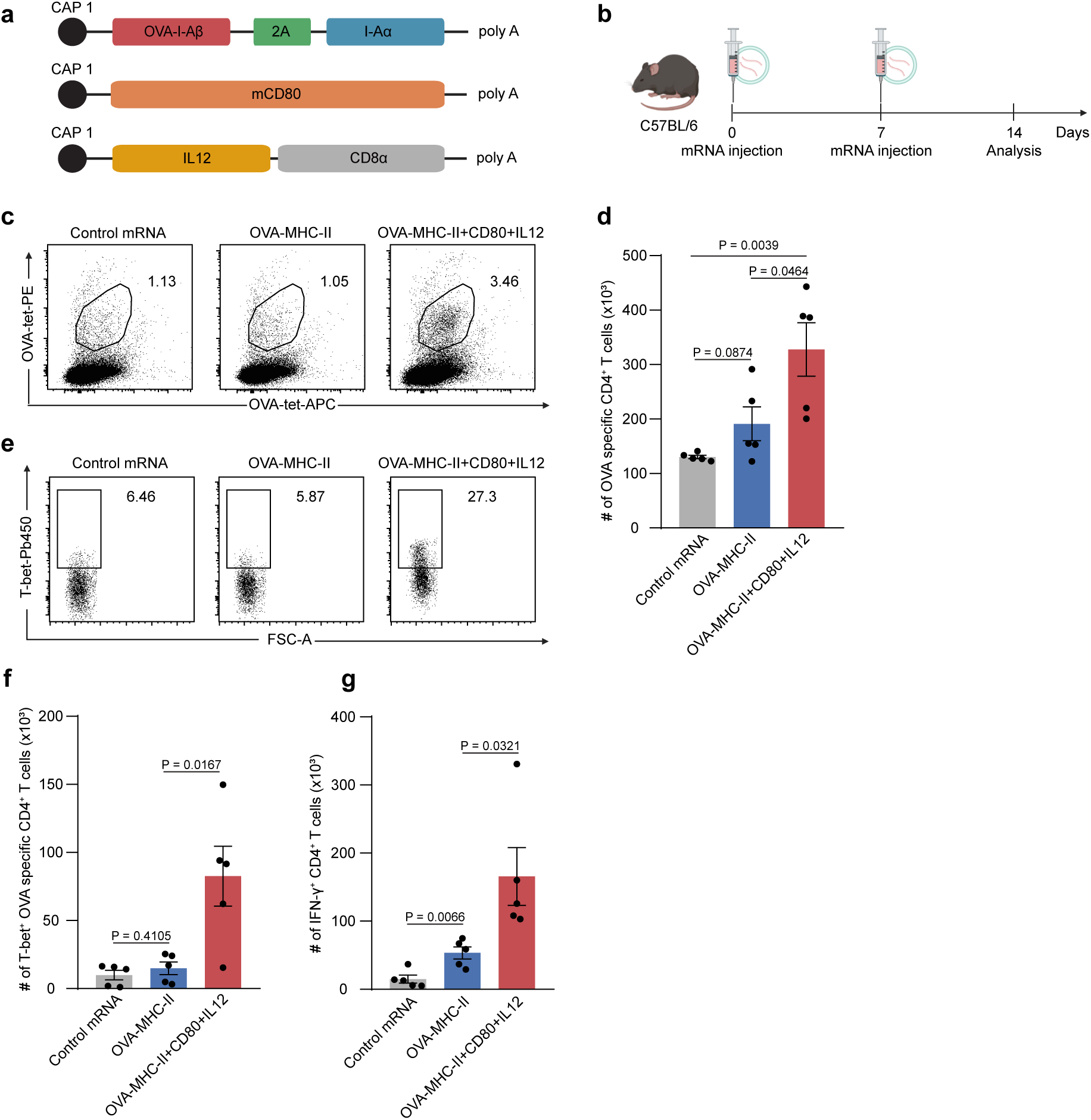
In vivo generation of designer APCs induces Th1 differentiation of antigen-specific CD4⁺ T cells. (a) Schematic representation of the mRNA construct encoding OVA-MHC-II, CD80 and membrane-anchored IL-12. (b) C57BL/6 mice received two weekly i.v. injections of control mRNA (15 µg), OVA-MHC-II mRNA (5 µg) plus control mRNA (10 µg), or OVA-MHC-II (5 µg) + CD80 (5 µg) + membrane-anchored-IL-12 (5 µg) mRNAs. The spleens were collected 7 d after the final dose. (c) Representative dot plots of OVA-specific CD4⁺ T cells in the spleens. (d) Number of tetramer⁺ CD4⁺ T cells in the spleens. (e) Representative dot plots of T-bet⁺ OVA-specific CD4⁺ T cells in the spleens. (f) Number of T-bet⁺ OVA-specific CD4⁺ T cells in the spleens. (g) Number of IFN-γ⁺ CD4⁺ T cells in the spleens. Data are mean ± SEM. Statistical significance for panels (d,f,g) was determined by using an unpaired, two-tailed Student’s t-test.

## Discussion

In this study, we demonstrated that co-delivery of mRNAs encoding antigens, co-stimulatory molecules, and immunomodulatory cytokines can transiently reprogram non-professional somatic cells into designer APCs in vivo, thereby markedly enhancing neoantigen-specific CD8⁺ T cell activation and expansion, overcoming the low immunogenicity and neoantigen-prediction challenges of conventional mRNA cancer vaccines. Furthermore, our results demonstrate that the cytokine composition of the mRNA mixture can be tuned to direct CD4⁺ T-cell differentiation. For example, inclusion of IL-12 selectively promoted Th1 polarization of antigen-specific CD4⁺ T cells. Several recent reports have examined IL-12 mRNA as a stand-alone adjuvant. Aunins *et al.* showed that formulating a separate IL-12 mRNA-LNP with an antigen-encoding mRNA-LNP boosts CD8⁺ T cell expansion and effector differentiation, but CD4⁺ T cell help remained modest^21^. Lapuente *et al.* achieved stronger Th1 skewing by anchoring membrane-tethered IL-12 in an adjuvant mRNA construct, yet the effect was still dependent on endogenous dendritic cells for optimal priming^22^. Unlike two-vial IL-12 mRNA adjuvant protocols, the designer APC mRNA cocktail, comprising antigen, CD80, and membrane-bound IL-12 effectively induced the in vivo expansion and Th1 polarization of endogenous OVA-specific CD4⁺ T cells, as evidenced by T-bet expression and IFN-γ secretion. These findings highlight the efficiency and modularity of the designer APC platform in directing antigen-specific helper T cell responses.

Beyond IL-12, incorporating IL-15 or IL-21 could potentiate memory-precursor CD8⁺ T cells^24, 25^, whereas co-expression of co-stimulatory molecules such as 4-1BBL or OX40L may further amplify effector functions^25, 26^. These observations underscore the platform’s modularity: tailored combinations of antigens, co-stimulatory molecules and cytokines can fine-tune T cell responses to specific needs. Beyond cancer vaccines, the approach could be adapted to induce antigen-specific tolerance in autoimmune or allergic disorders by co-encoding inhibitory cytokines or other regulatory cues^28, 29^. Other applications, such as the induction of long-lived memory T cells or the avoidance of T cell exhaustion, may also be explored by further customizing the mRNA payload. Thus, designer APCs offer a flexible and precise platform for immune modulation in a variety of disease contexts.

In this study, we employed commercially available liposomes for mRNA delivery, which efficiently facilitated uptake and expression in recipient cells. However, the use of more advanced delivery vehicles, such as LNPs or LPX engineered for targeted delivery to dendritic cells, may further enhance transfection efficiency and therapeutic outcomes. Recent reports have shown that dendritic-cell-targeting LPX, such as those developed by BioNTech (e.g., BNT111) and others, can improve antigen presentation and anti-tumor immunity^14, 29, 30^. Beyond lipid-based carriers, viral-like particles, polymer-lipid hybrid nanoparticles, and engineered extracellular vesicles represent additional platforms with the potential to increase cargo stability, cellular specificity, and endosomal escape^31–33^. Integration of the designer APC mRNA platform with advanced delivery systems may further enhance anti-tumor immunity by improving the precision and efficiency of mRNA delivery to immune-relevant cell populations.

While these advantages are compelling, the potent immune activation achieved with designer APCs raises safety considerations. Excessive or systemic expression of pro-inflammatory cytokines could precipitate cytokine-release syndrome or off-target tissue damage^34, 35^. Rigorous dose-finding studies, and tissue-restricted delivery will therefore be critical components of future translational efforts.

In summary, our study establishes a novel and modular mRNA-based platform for the in vivo generation of designer APCs, enabling the precise activation and programming of antigen-specific T cell responses. This work provides a foundation for future investigations into optimized mRNA formulations, advanced delivery systems, safety engineering, and translational applications in cancer immunotherapy and immune regulation.

## Methods

### Cell lines

The OVA-expressing murine lymphoma cell line, a derivative of EL4 (E.G7; ATCC, CRL-2113) was cultured in RPMI 1640 (Nacalai Tesque) supplemented with 10% fetal calf serum (FCS; Thermo Fisher Scientific), 1× non-essential amino acids (Nacalai Tesque), 1 mM sodium pyruvate (Nacalai Tesque), 100 U/mL penicillin, 100 U/mL streptomycin (FUJIFILM Wako), and 0.05 μM 2-mercaptoethanol (Thermo Fisher Scientific). To generate a stable MC38 cell line expressing mutant Rpl18, we used a codon-optimized mouse Rpl18 gene containing the Q125R mutation, which was previously reported in MC38 cells^36^, but absent in the MC38 cells obtained from Kerafast. The mutated gene was fused to the transmembrane domain of mouse CD8 via a 2A peptide and cloned into a third-generation lentiviral vector (pLJM1, Addgene #91980) under the CMV promoter. MC38 cells were transduced with lentiviral particles and stably expressed mutant Rpl18. Rpl18-MC38 cells were cultured in DMEM (Nacalai Tesque) supplemented with 10% FCS, 2 mM L-glutamine (Nacalai Tesque), 1× non-essential amino acids, 1 mM sodium pyruvate, 10 mM HEPES (Nacalai Tesque), 50 μg/mL gentamicin sulfate (Nacalai Tesque), and 100 U/mL penicillin-streptomycin. Human embryonic kidney (HEK)-293T cells (ATCC, Cat# CRL-3216) were cultured in DMEM supplemented with 10% FCS, 100 U/mL penicillin, and 100 U/mL streptomycin. All cells were maintained at 37 °C in a humidified atmosphere containing 5% CO₂.

### Mice

C57BL/6 mice were purchased from Japan SLC. OT-I TCR transgenic mice^37^ and HLA-A*02:01 transgenic mice^18^ mice were housed in a specific pathogen-free facility. All animal experiments were performed following a protocol approved by Kanazawa University.

### mRNA construct and in vitro transcription

Plasmids for mRNA preparation were codon-optimized for murine expression using VectorBuilder (VectorBuilder Ltd.). Each construct contained a consensus Kozak sequence upstream of the start codon and three stop codons downstream of the gene of interest. DNA fragments were synthesized by Eurofins Genomics and cloned into the vector, which contains a T7 RNA polymerase promoter, 5′UTR, coding region, 3′UTR, and a 128-base poly(A) tail. The 5′UTR, coding region, and 3′UTR sequences were based on those described in a previous study^38^.

In vitro transcription was performed using the HiScribe® T7 High Yield RNA Synthesis Kit (New England Biolabs). Transcribed mRNA was purified using the Monarch® RNA Cleanup Kit (New England Biolabs). The quality of IVT-mRNA was confirmed with the RNA ScreenTape kit (Agilent Technologies).

### mRNA transfection in vitro

mRNA transfections were performed using the TransIT®-mRNA Transfection Kit (Mirus Bio), a cationic lipid-based reagent. Briefly, mRNA was diluted in Opti-MEM (Thermo Fisher Scientific) and mixed with TransIT-mRNA reagent according to the manufacturer’s protocol. The resulting RNA-lipid complexes were added to the cells. Target protein expression was evaluated by flow cytometry 18 hours after transfection.

### In vitro T cells proliferation assay

Lymph node T cells were isolated from OT-I transgenic mice and labeled with 1 μM CellTrace™ Violet (CTV; Thermo Fisher Scientific) at 37 °C for 3 minutes. A total of 2 × 10⁵ CTV-labeled OT-I T cells were co-cultured with MC38 cells transfected with mRNA encoding OVA-MITD, OVA-MITD + CD80, OVA-MITD + membrane-anchored IL-2, or OVA-MITD + CD80 + membrane-anchored IL-2 at a 4:1 T cell to MC38 cell ratio. Transfected MC38 cells were prepared according to the mRNA transfection method described above. After 3 days of culture, T cell proliferation was analyzed by flow cytometry.

### mRNA formulation and intravenous administration

mRNA was formulated with in vivo-jetRNA+ (Polyplus) according to the manufacturer’s instructions. Briefly, mRNA was diluted in a suitable buffer and mixed with in vivo-jetRNA+ reagent. A total of 200 μL of the resulting nanoparticle suspension was intravenously injected into each mouse.

### In vivo tumor models and mRNA treatment regimen

A total of 1 × 10⁵ E.G7 tumor cells suspended in 200 μL PBS (FUJIFILM Wako) were subcutaneously injected into the right flank of C57BL/6 mice. For the Rpl18-MC38 tumor model, 5 × 10⁵ Rpl18-MC38 cells suspended in a 1:1 mixture of 100 μL sterile PBS and 100 μL Matrigel (Corning) were injected into the same site. For both tumor models, when tumors reached approximately 100 mm³ in volume, mice received weekly intravenous injections of mRNA for three consecutive weeks. Treatment consisted of either OVA-MITD (5 μg) or Rpl18-MITD (5 μg) combined with control mRNA (5 μg), or a mixture of OVA-MITD (5 μg) or Rpl18-MITD (5 μg), CD80 mRNA (5 μg), and membrane-anchored IL-2 mRNA (5 μg). As a negative control, control mRNA (15 μg) alone was administered. Akaluc-Venus mRNA was used as the control mRNA in all treatment groups^39^. Tumor volumes were measured two to three times per week using digital calipers and calculated using the formula (length × width²) / 2, where length and width represent the longest and shortest tumor diameters, respectively. Mice were euthanized when tumors reached ≥2000 mm³, exhibited ulceration, or showed signs of severe distress or moribundity, in accordance with predefined humane endpoint criteria.

### Isolation and Preparation of Tumor-Infiltrating Lymphocytes (TILs)

Tumors were harvested and minced into ∼2-3 mm fragments. Tissue dissociation was performed using the Tumor Dissociation Kit, mouse (Miltenyi Biotec) following the manufacturer’s protocol. Briefly, tumor fragments were incubated with enzyme mix in RPMI 1640 medium and processed on the GentleMACS™ Octo Dissociator (Miltenyi Biotec). The resulting single-cell suspension was filtered through a 70 μm strainer to remove debris. Cells were washed twice with PBS containing 2% FCS and subsequently counted. Purified TILs were resuspended in complete RPMI 1640 medium containing 10% FCS for subsequent flow cytometric analysis.

### TCR transduction of primary human T cells and co-culture experiment

Peripheral blood mononuclear cells (PBMCs) from HLA-A*02:01-positive healthy donors were cultured in X-VIVO 15 medium (Lonza) containing 5ng/mL human IL-2 (BioLegend) and stimulated using Dynabeads Human T-Activator CD3/CD28 (Thermo Fisher Scientific). After 48 hours, the beads were removed, and the activated T cells were transduced with a lentiviral vector encoding a TCR specific for the NY-ESO-1 peptide (amino acids 157-165, SLLMWITQC)^17^. The medium was subsequently refreshed three days after transduction. NY-ESO-1 TCR-engineered T cells were labeled with 1μM CTV at 37 °C for 3 minutes. Subsequently, 2 × 10⁵ CTV-labeled T cells were co-cultured with HEK293T cells transfected with either control mRNA, NY-ESO-1-MITD mRNA alone, or NY-ESO-1-MITD combined with hCD80 and membrane-anchored hIL-2 mRNAs, at a T cell to HEK293T cell ratio of 4:1. CD69 expression was measured after 24 hours of co-culture, while IFN-γ production and T cell proliferation were evaluated by flow cytometry three days after co-culture initiation.

### In vivo killing assay

HLA-A*02:01 transgenic (HHD) mice were intravenously injected with in vivo-jetRNA® complexes containing either a designer APC cocktail of NY-ESO-1-MITD (5 µg), murine CD80 (5 µg), and membrane-anchored murine IL-2-CD8 (5 µg); NY-ESO-1-MITD alone (5 µg); or Akaluc-Venus (15 µg) as a negative control. Seven days later, 4 × 10⁶ syngeneic target splenocytes, prepared as a 1:1 mixture of NY-ESO-1-pulsed cells labeled with CTV and Wilms Tumor 1 peptide-pulsed cells labeled with CFSE (Thermo Fisher Scientific), were transferred intravenously. For peptide loading, HLA-A02:01 splenocytes (1× 10⁷ cells /ml) were incubated with 10 µmol peptide for 2 h at 37 °C and washed twice by centrifugation to remove unbound peptide. Twenty hours after target-cell transfer, spleens were harvested, converted to single-cell suspensions, and analyzed by flow cytometry; antigen-specific cytotoxicity was quantified from the preferential loss of CTV⁺ NY-ESO-1-pulsed cells relative to CFSE⁺ control peptide-pulsed cells.

### Antibodies and Flow Cytometry

Antibody staining was performed according to standard procedures. Monoclonal antibodies for surface staining, including antibodies against mCD3 (clone: 17A2), mCD4 (clone: GK1.5), mCD8α (clone: 53-6.7), mCD44 (clone: IM7), mCD45 (clone: 30-F11), mCD62L (clone: MEL-14), mCD80 (clone: 16-10A1), mTCR-Vα2 (clone: B20.1), mH-2K^b^ bound to SIINFEKL-PE (clone: 25-D1.16), mIL-2 (clone: JES6-5H4), hCD4 (clone: RPA-T4), hCD8 (clone: SK1), hCD69 (clone: FN50) were purchased from BioLegend. Intracellular staining was performed using antibodies against mIFN-γ (clone: XMG1.2), hIFN-γ (clone: 4S.B3), mT-bet (clone:4B.10) in combination with the True-Nuclear Transcription Factor Buffer Set (BioLegend). OVA-specific CD8⁺ T cells, Rpl18-specific CD8⁺ T cells, NY-ESO-1-specific CD8⁺ T cells, and OVA-specific CD4⁺ T cells were quantified using the following tetramers obtained from the NIH Tetramer Core Facility: H-2Kᵇ/OVA (257-264), H-2Kᵇ/Rpl18 (KILTFDRL), HLA-A*02:01-H2Kᵇ/NY-ESO-1 (SLLMWITQC), and I-Aᵇ/OVA (323-339), respectively. Staining was performed according to the manufacturer’s instructions. For intracellular staining, cells were fixed and permeabilized using the True-Nuclear Transcription Factor Buffer Set (BioLegend) according to the manufacturer’s instructions. Briefly, cells were incubated with fixation buffer for 60 minutes at room temperature in the dark, followed by two washes with permeabilization buffer. Cells were then stained with antibodies diluted 1:50 in permeabilization buffer for 45 minutes at room temperature in the dark. After washing, cells were resuspended in FACS buffer and subjected to flow cytometric analysis. Data were acquired using FACSDiva software (BD). A FACSMelody (BD) was used for cell sorting. Data analysis was performed using FlowJo v.10.4.1 software.

## ACKNOWLEDGMENTS

This work was supported by World Premier International Research Center Initiative (WPI), MEXT, Japan. We thank T. Yoshida for the helpful discussion. We thank the NIH Tetramer Core Facility for providing H-2Kᵇ/OVA (257-264), H-2Kᵇ/Rpl18 (KILTFDRL), HLA-A*02:01-H2Kᵇ/NY-ESO-1 (SLLMWITQC), and I-Aᵇ/OVA (323-339) tetramers. This work was supported by World Premier International Research Center Initiative (WPI), MEXT, Japan, Doctoral Program for World-leading Innovative & Smart Education (WISE) Program for Nano-Precision Medicine (TVL, IS), Japan Science and Technology Agency (JST) Fusion Oriented Research for Disruptive Science and Technology (FOREST) grant no. JPMJFR2115 (TY), and Practical Research for Innovative Cancer Control from the Japan Agency for Medical Research and Development (AMED) grant no. 24ck0106967h0001 (TY). ChatGPT (GPT-4, OpenAI) was used to refine the language and enhance readability. The authors take full responsibility for the final content.

## Data availability

Upon reasonable request, data can be obtained from the corresponding author.

## Author Contributions

T.V.L.: Conceptualization, Investigation, Data curation, Formal analysis, Visualization, Writing-original draft. S.I., I.F., M.U.: Investigation, Methodology, Validation, Visualization. S.N., S.M.: Resources, Methodology. R.H.: Conceptualization, Supervision, Funding acquisition, Writing-review & editing. T.Y.: Conceptualization, Methodology, Supervision, Funding acquisition, Writing-review & editing.

## Ethics approval statement

All animal experiments were conducted in compliance with the ARRIVE guidelines (https://arriveguidelines.org/) and approved by the Animal Experimentation Committee of Kanazawa University (Kanazawa, Ishikawa, Japan; approval number: AP-204131). Female C57BL/6 mice aged 6 to 8 weeks and weighing 18-22 g were used. A total of 120 animals were included. Mice were group-housed (up to five per cage) under specific pathogen-free conditions, with ad libitum access to standard rodent chow and water, and maintained in a controlled environment (temperature, humidity, and 12-hour light/dark cycle). Environmental enrichment (e.g., nesting materials and shelters) was provided to minimize stress. At the end of the experiments, animals were euthanized using CO₂ inhalation with a gradual fill rate, in accordance with AVMA guidelines. PBMCs were collected with written informed consent from single healthy volunteers following the guidelines approved by the Research Ethics Committee of Kanazawa University.

---

**Supplementary Figure 1.**
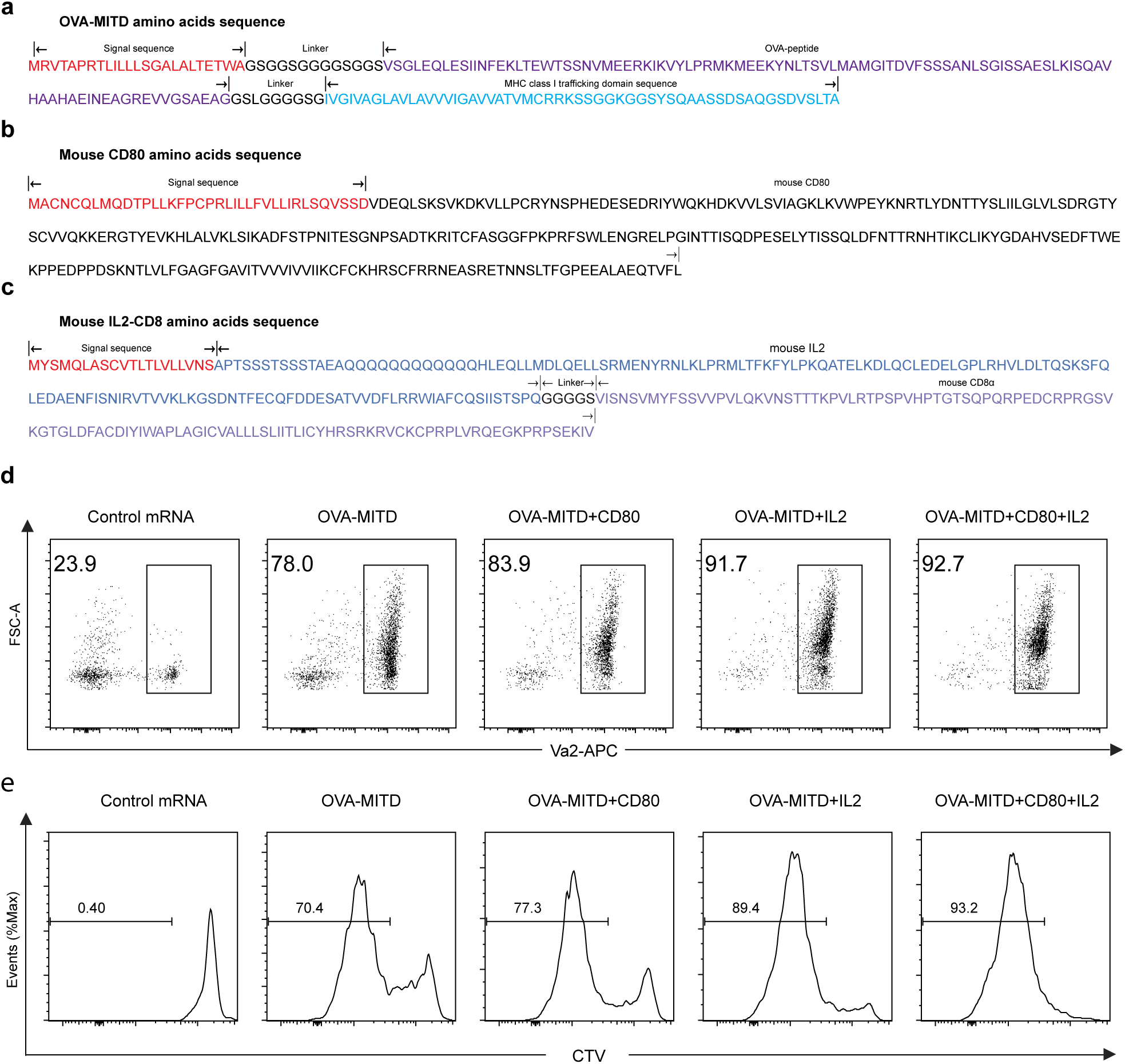
Designer APC-mediated stimulation of CD8⁺ T cells. (a-c) Amino acid sequences of OVA-MITD, mouse CD80, and membrane-anchored mouse IL-2 used for mRNA synthesis. (d, e) Representative flow cytometry plots showing OT-I CD8⁺ T cells and proliferating (CTV^low^) OT-I cells after co-culture with designer APCs.

**Supplementary Figure 2.**
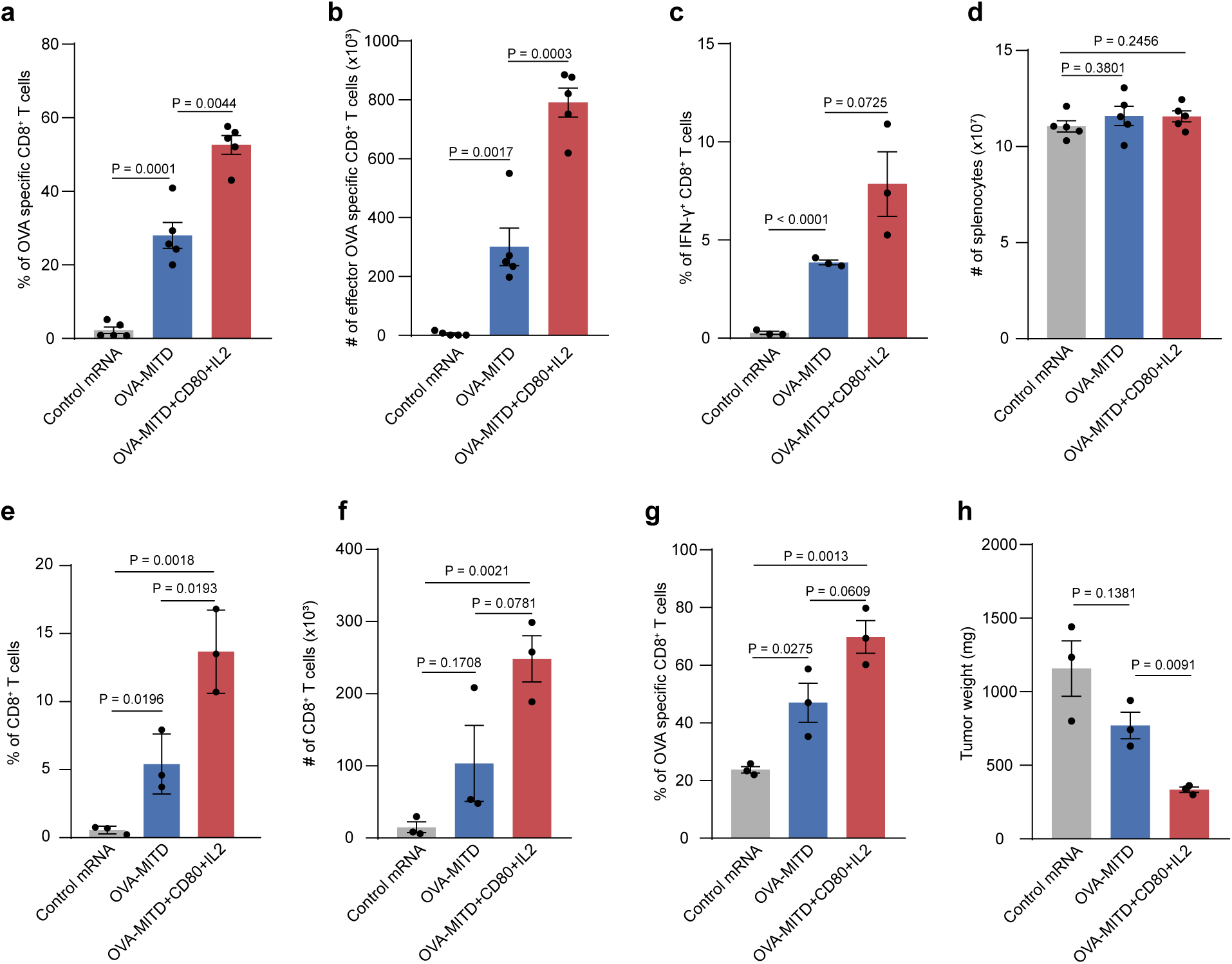
Designer APC-induced activation of antigen-specific CD8⁺ T Cells in spleen and tumor. (a) Frequencies of OVA-specific CD8⁺ T cells in the spleens. (b) Number of CD44^hi^CD62L^low^ CD8⁺ T cells in the spleens. (c) Frequencies of IFN-γ⁺ CD8⁺ T cells in the spleens. (d) Number of total splenocytes. (e) Frequencies of CD8⁺ T cells in the tumors. (f) Quantification of CD8⁺ T cells per 100 mg tumors. (g) Frequencies of OVA-specific CD8⁺ T cells in TILs. (h) Tumor weight at harvest. Data are presented as mean ± SEM. Statistical significance was assessed using unpaired, two-tailed Student’s *t*-test.

**Supplementary Figure 3.**
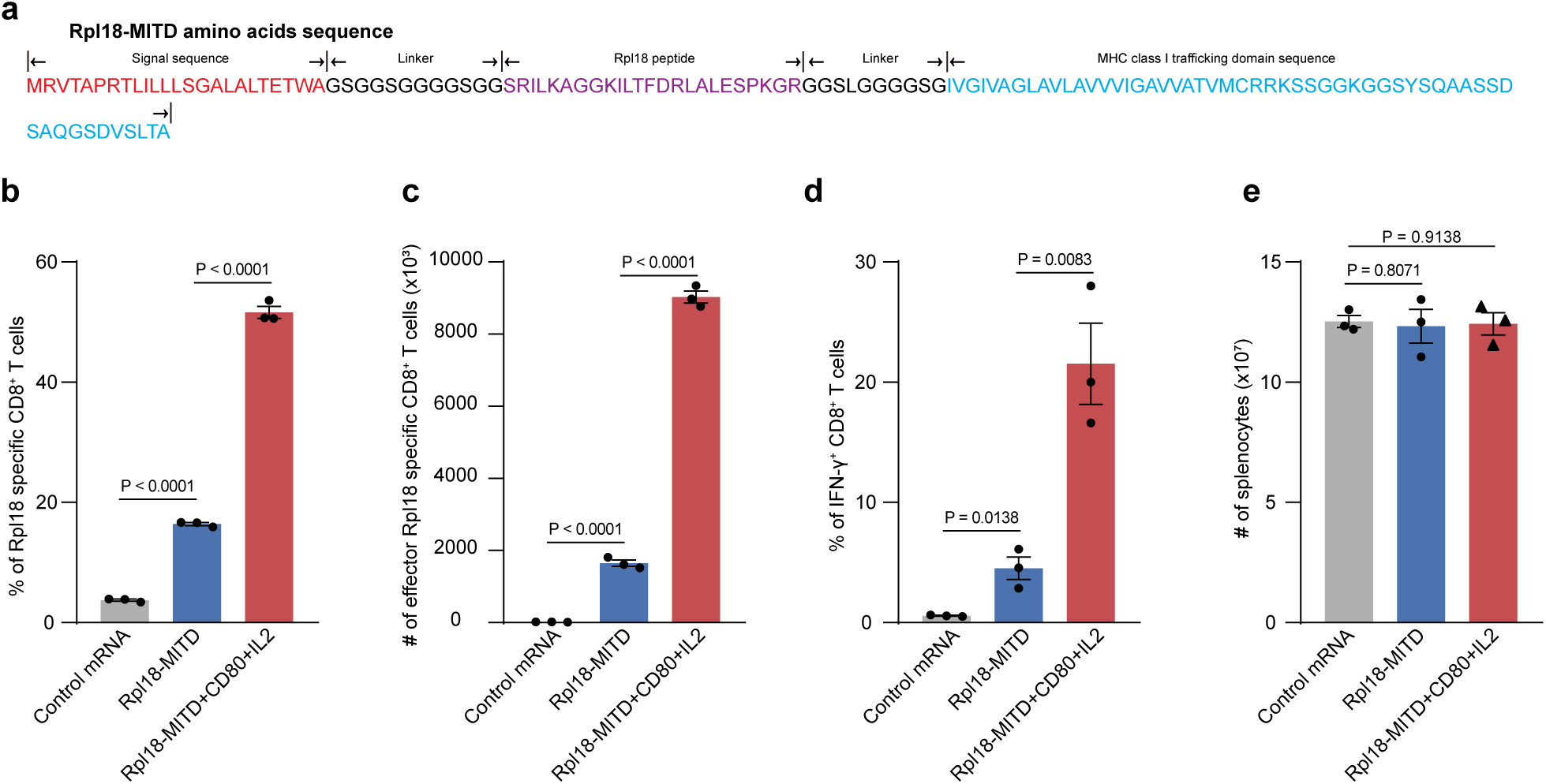
Activation of neoantigen-specific CD8⁺ T cells by designer APCs. (a) Amino acid sequence of Rpl18-MITD. (b) Frequencies of Rpl18-specific CD8⁺ T cells in the spleens. (c) Number of CD4^hi^CD62L^low^ CD8⁺ T cell in the spleens. (d) Frequencies of IFN-γ⁺ CD8⁺ T cells in the spleens. (e) Number of total splenocytes. Data are presented as mean ± SEM. Statistical significance was assessed using unpaired, two-tailed Student’s *t*-test.

**Supplementary Figure 4.**
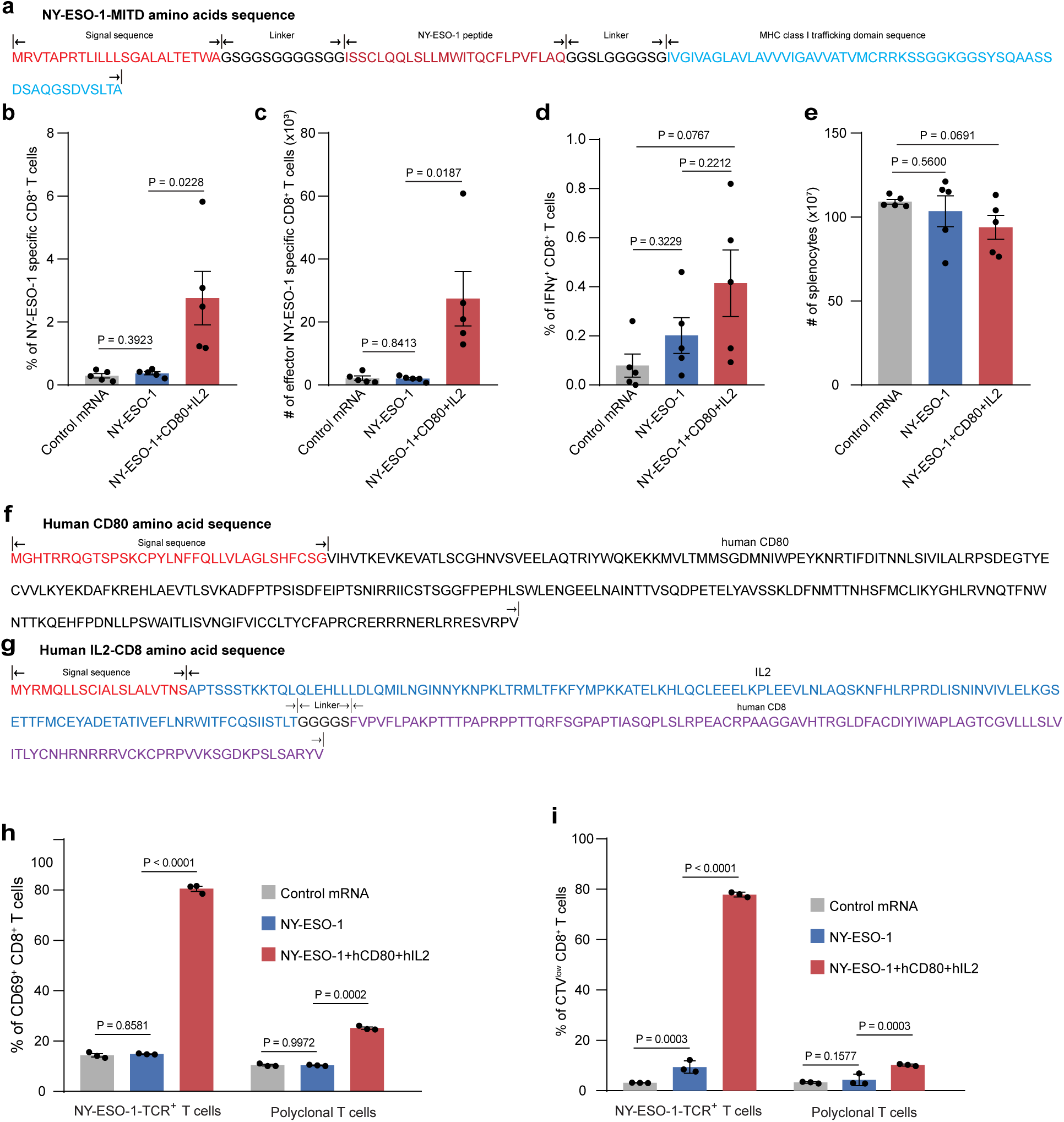
Humanized designer APCs activate antigen-specific CD8⁺ T cells. (a) Amino acid sequence of NY-ESO-1-MITD. (b) Frequencies of NY-ESO-1 specific CD8⁺ T cells. (c) Number of CD44^hi^CD62L^low^ NY-ESO-1 specific CD8⁺ T cell in the spleens. (d) Frequencies of IFN-γ⁺ CD8⁺ T cells in the spleens. (e) Number of total splenocytes. (f) Amino acid sequence of human CD80. (g) Amino acid sequence of membrane-anchored human IL-2. (h) Frequencies of CD69^+^ CD8 T cells. (i) Frequencies of CTV^low^ CD8⁺ T cells. Data are presented as mean ± SEM. Statistical significance was determined using an unpaired, two-tailed Student’s t-test for panels (b-e), and two-way ANOVA followed by Tukey’s multiple comparisons test for panels (h) and (i).

**Supplementary Figure 5.**
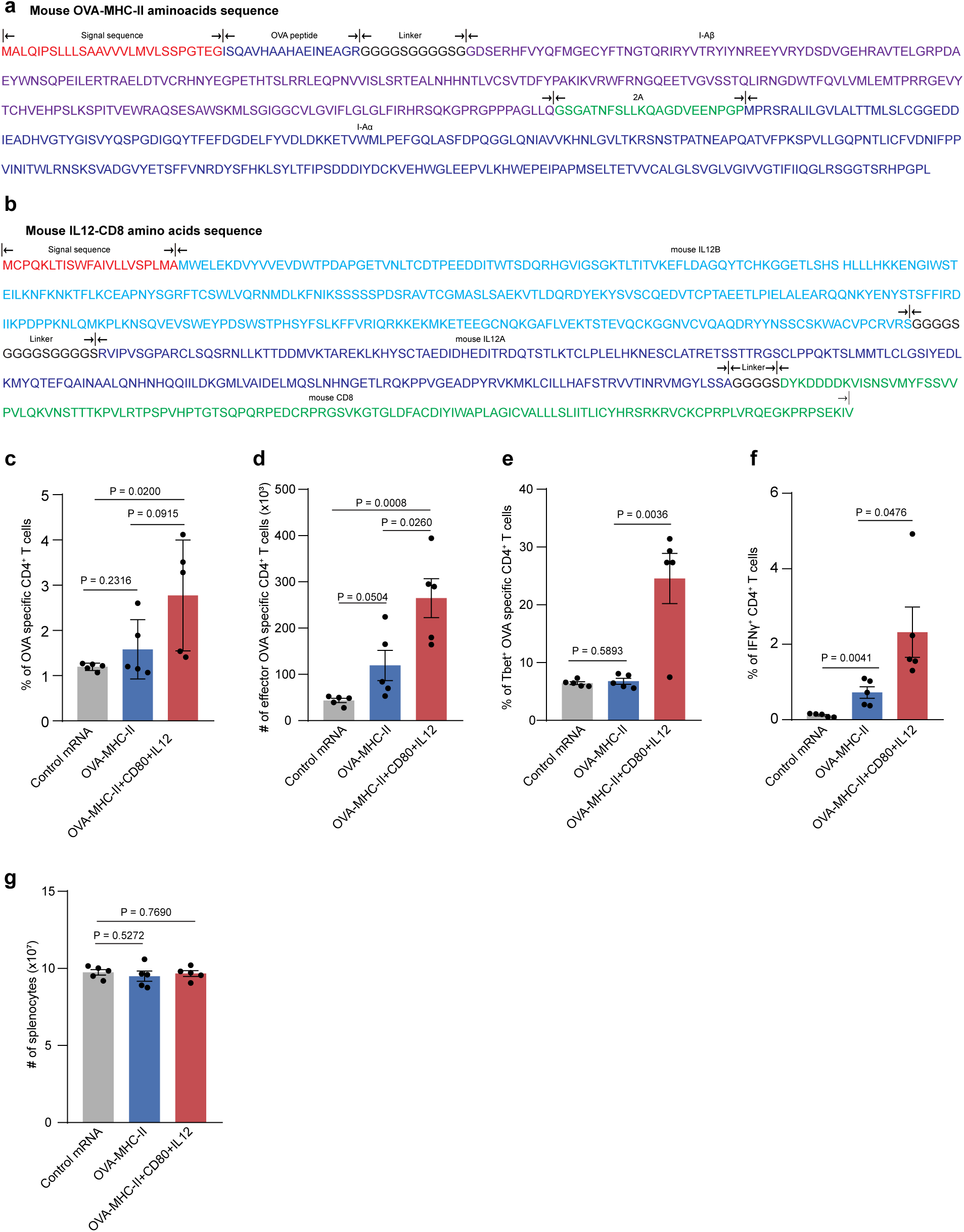
Induction of Th1 cells by designer APCs. (a) Amino acid sequence of membrane-anchored mouse IL-12. (b) Schematic of OVA peptide fused to MHC-II. (c) Frequencies of OVA-specific CD4⁺ T cells. (d) Number of CD44^hi^CD62L^low^ CD4⁺ T cell phenotype. (e) Frequencies of T-bet⁺ CD4⁺ T cells. (f) Frequencies of IFN-γ⁺ OVA-specific CD4⁺ T cells. (g) Number of total splenocytes. Data are presented as mean ± SEM. Statistical significance was assessed using unpaired, two-tailed Student’s *t*-test.

